# Pdl1 and icosl discriminate human secretory and helper dendritic cells

**DOI:** 10.1101/721563

**Authors:** Caroline Hoffmann, Floriane Noel, Maximilien Grandclaudon, Paula Michea, Aurore Surun, Lilith Faucheux, Philemon Sirven, Olivier Lantz, Juliette Rochefort, Jerzy Klijanienko, Charlotte Lecerf, Maud Kamal, Christophe Le Tourneau, Maude Guillot-Delost, Vassili Soumelis

## Abstract

Dendritic cells (DC) are described as immature at the steady state, with a high antigen capture capacity, turning into a mature state with a strong T cell stimulatory capacity upon activation. Using 16 different stimuli in vitro (130 observations), we describe two states of human activated dendritic cells. PDL1highICOSLlow “secretory DC” produced large amounts of inflammatory cytokines and chemokines but induced very low levels of T helper (Th) cytokines following DC-T co-culture; conversely PDL1lowICOSLhigh “helper DC” produced low levels of secreted factors but induced high levels of Th cytokines characteristic of a broad range of Th subsets. Secretory DC were phenotypically identified in T cell inflamed primary head and neck squamous cell carcinoma. RNAseq analysis showed that they expressed a typical secretory DC signature, including CD40, PVR, IL1B, TNF, and CCL19. This novel and universal functional dichotomy of human DC opens broad perspectives for the characterization of inflammatory diseases, and for immunotherapy.

## MANUSCRIPT

Dendritic cells (DC) have a key role in initiating and polarizing the immune responses, including anti-tumor immunity (1). Immature DC patrol in tissues and have a low expression of costimulatory molecules. Following antigen stimulation, they mature and acquire a strong T cell stimulatory capacity (2). So far, mature DC have been classified as immunogenic when they induced T effectors and secreted IL12 and IL1b, or tolerogenic when they induced regulatory T cells and secreted IL-10, TNF and TGFb, or no cytokines (3), (4), (5), (6). In cancer, it is considered that factors derived from the tumor microenvironment induce tolerogenic DC (7), (8), (9). However, most studies have been realized using a limited number of stimuli, mostly in mice models or in vitro with human monocyte-derived DC (10), (11), (12). Furthermore, the phenotype and function of tissue infiltrating DC in human remains largely unknown. Our aim was to decipher the mechanisms regulating DC phenotypes and to understand their associated function, with a physiopathological relevance in human cancer.

To determine the phenotypic heterogeneity of DC infiltrating cancer tissue and its relation to the other immune cell types, we analyzed by flow cytometry 22 fresh head and neck squamous cell carcinoma (HNSCC) samples. Here, we show that the frequencies of tumor infiltrating CD3 T cells were positively associated to the frequencies of DC and to PDL1 expression on CD11c+HLA-DR+ cell subsets, and negatively associated to the frequencies of neutrophils and of ICOSL expression on the same cells (Fig1). We used 2 different antibody panels analyzing T cell subsets (Fig S1A) and myeloid cells subsets (Fig 1A, 1B). In the myeloid panel, CD45+, Lineage− (CD3, CD19, CD56) cells were analyzed by their expression of CD11c and HLA-DR. The double positive population was separated into four populations by their expression of CD14 and BDCA1, and included the monocytes and macrophages (MMAC), the CD14+DC, the cDC2 (BDCA1+CD14-) and the double negative population enriched in cDC1 (cDC1e) (Fig S1B). Plasmacytoid DC were gated as CD11c-, HLA-DR+, CD123+. We extracted a total of 434 parameters. We found a large variation of CD3 infiltration across tumors ranking from 1% to 61% (Fig 1C). In order to identify the parameters associated to tumor inflammation, we defined 3 groups of equivalent sizes labeled “CD3 High” (n=8), “CD3 Int” (n=6), and “CD3 low” (n=8). To avoid bias, we used a sub-list of 81 non-redundant parameters among the 434 measured, meaning that each population was expressed only in percentage of its parental population (Table 1). CD3 high tumors were significantly enriched in cDC2, cDC1e, pDC and in PDL1 expressing MMAC and cDC1e. Conversely, CD3 low tumors were enriched in Lin-DR-cells (mainly neutrophils, see Fig S1C), macrophages, and ICOSL expressing CD11c+HLA-DR+ cells (Fig 1D, 1E, 1F). The levels of expression of PDL1 and ICOSL in the four CD11c+HLA-DR+ subsets were highly correlated in all tumor samples (Fig S1D). CD11c+HLA-DR+ cell subsets in CD3 low tumors expressed intermediate levels of PDL1 and ICOSL, and were closer to the expression observed on their blood counterparts than the same subsets in CD3 high tumors, which upregulated PDL1 and downregulated ICOSL (Fig 1E). Thirteen out of the 16 significant parameters were obtained from the myeloid cell panel (Fig 1D), showing that there were fewer variations in the percentages of the various T cells subsets related to CD3 infiltration levels. For example, the proportion of regulatory T cells among the CD4+ T cells were 34%, 35% and 41% in the CD3 High, Int and Low groups respectively. Finally, to determine if any combined parameter, ratio or clinical variable was highly efficient at discriminating the 3 groups, we performed an elastic net model including the all the 434 parameters and 14 clinical parameters (Table 2). We found that the intermediate expression of ICOSL on CD11c+HLA-DR+ cell subsets was highly characteristic of the CD3 low group (Fig S1E). Only parameters directly linked to T cell infiltration (percentages of T cell subsets in live cells) were found in the high CD3 group. In summary, we showed that CD3 inflamed tumors were more infiltrated by DC subsets that expressed higher levels of PDL1 than in non-inflamed tumors, and that PDL1 and ICOSL expressions on DC and macrophages were opposed (Fig 1D, Fig S1D).

**Fig1.**
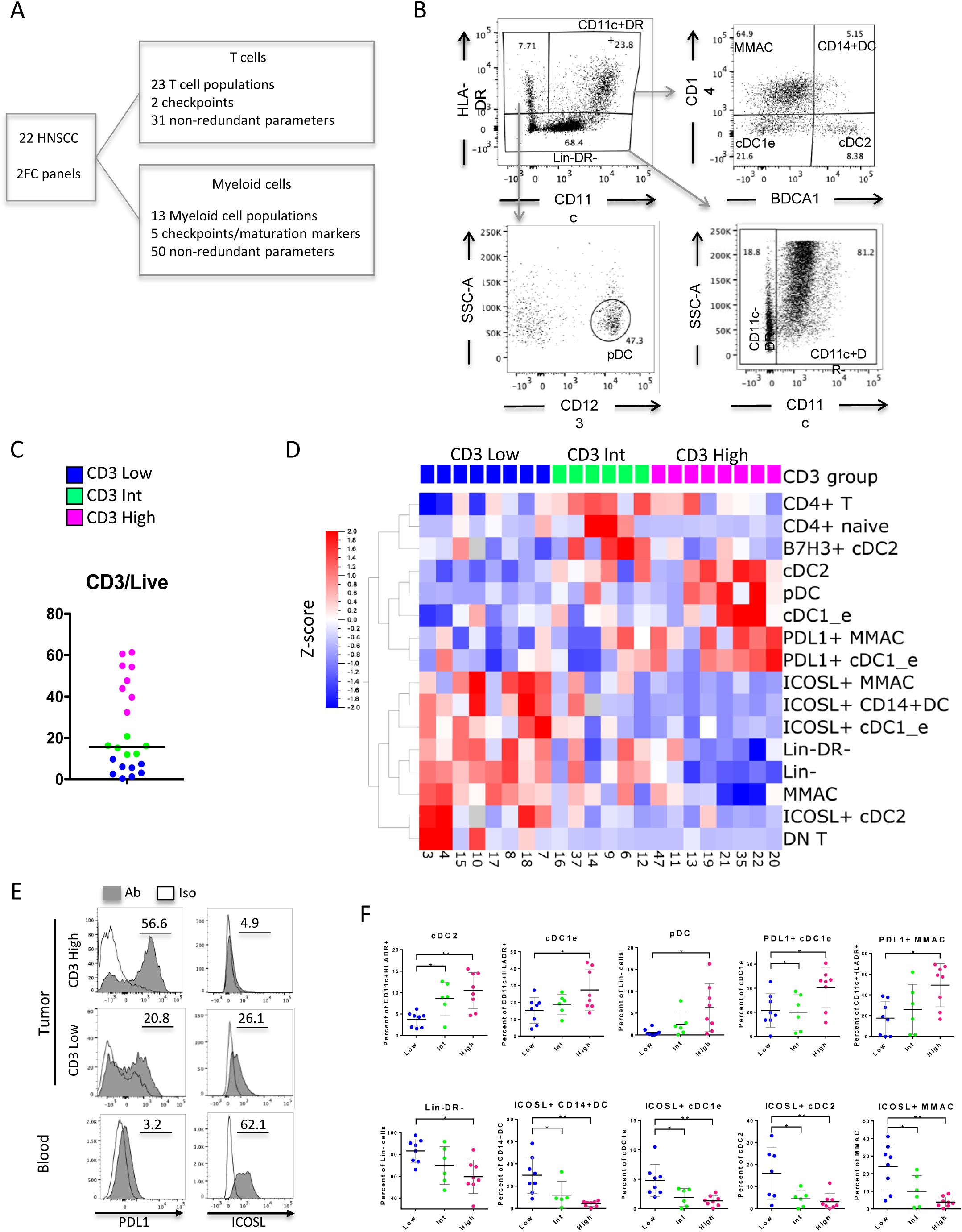
T cell infiltration is associated to DC infiltration and PDL1 & ICOSL expression on CD11c+HLA-DR+ cell. Phenotypic characterization of 22 human HNSCC primary tumor-infiltrating cells. A. Multicolor flow cytometry analysis scheme. B. Myeloid cell panel gating strategy for the CD45+CD3-CD16-CD19-compartment. C. Percentage of CD3 positive cells among live cells. D. Anova test between CD3 high, int and low, showing only the 16 significant variables among the 81 analyzed. E. Representative staining of PDL1 (right) and ICOSL (left) in CD11c+DR+ cells in a representative CD3 high tumor (top), CD3 low (middle) and blood from a healthy donor (bottom). F. Quantification of cell populations in percentages of their parental population in the 3 groups of CD3 infiltration.

To identify candidate stimuli that could be responsible for the PDL1/ICOSL expression patterns and to further understand the subsequent functional implications, we took advantage of a DC-T cell dataset from Grandclaudon et al. (13). We used the existing data on primary blood CD11c+HLA-DR+ DC and generated supplementary experiments and analysis. Briefly, blood DC were activated for 24 hours by 16 different types of perturbators and analyzed for their expression of 29 surface markers (n=154 data points), and their secretion of 32 chemokines and cytokines (n=130 data points). The remaining cells were co-cultured with allogenic naïve CD4 T cells for 6 days and we measured the expansion fold. After 24h of restimulation by anti CD3/CD28 we measured 17 T helper cytokines (Fig 2A). We confirmed the anti-correlation of PDL1 and ICOSL expression (Fig2B). Three main groups of responses were observed: (i) PDL1high and ICOSLlow, like on ex vivo cDC2 from inflamed tumors; (ii) PDL1low and ICOSLhigh, and (iii) medium-like PDL1 low and ICOSL low (Fig 2B). Co-expression of both PDL1high and ICOSLhigh was a rare profile and was not observed for very high expression levels. ICOSL expression was null when PDL1 expression reached its highest levels. We used an unsupervised approach by t-SNE of the 29 surface markers to verify that PDL1 and ICOSL were relevant markers to discriminate the various DC phenotypes observed in vitro. We observed that PDL1 high cells clustered together and were distinct from ICOSL high cell clusters and from PDL1 low ICOSL low cluster, the latter including most Medium-DC conditions (Fig 2C). The DC perturbators inducing a majority of PDL1 high ICOSL low cDC2 were R848, Zymosan, HKSA and HKLM, while the ones inducing a majority of ICOSL high PDL1 low cDC2 were TSLP, GM-CSF and Flu (Fig 2C, Table 3). To pursue the analysis of the different functions of these DC phenotypes, we defined 4 groups of activated DC by their PDL1 and ICOSL expression (Fig 2B). First, we analyzed the 29 surface markers in these 4 groups and in Medium-DC: PDL1 High ICOSL low DC co-expressed PVR, PDL2, Nectin2, CD54, and CD40, with Spearman correlation coefficients of 0.8, 0.75, 0.66, 0.64 and 0.62 respectively (Fig2D, 3E, Table 4). ICOSL high PDL1 low DC did not have any correlated molecule with a Spearman correlation coefficient superior to 0,5.

**Fig 2.**
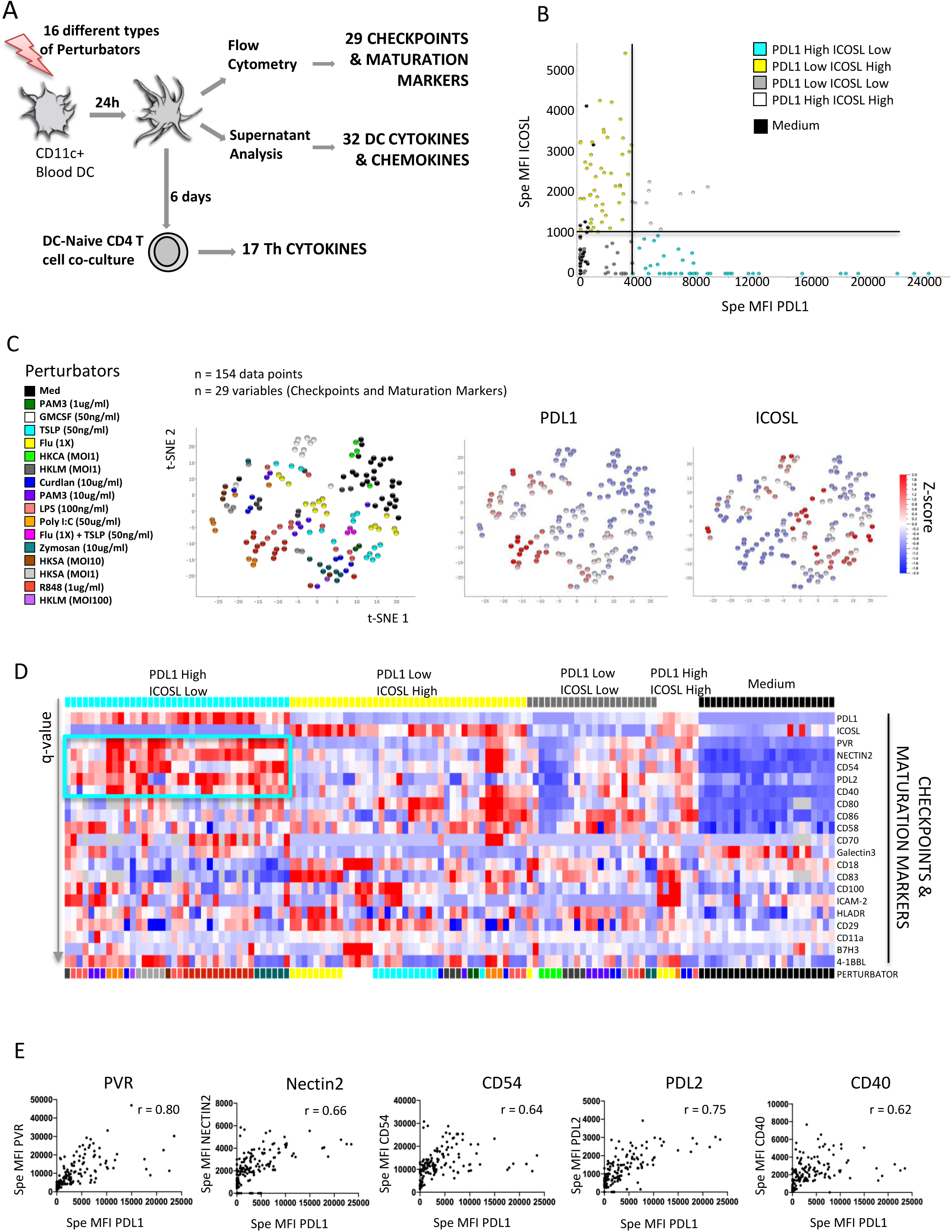
PDL1 and ICOSL expression on CD11c+DC were exclusive and PDL1 high DC overexpress PVR, Nectin2, CD54, CD40 and PDL2. A. Methods for the in vitro analysis of primary blood DC. B. Expression of PDL1(x) vs ICOSL(y) on DC at H24. Individual tests were annotated according to their expression of PDL1 as high/low and ICOSL high/low with the thresholds of specific MFI at 3500 and 1000 respectively. C. T-SNE of the 29 surface markers colored by stimuli (left), PDL1 specific MFI (center) and ICOSL specific MFI (right) using Qlucore software. D. Heatmap representing the expression of the 29 surface markers in the 4 groups defined by PDL1 and ICOSL in “B”, and in Medium condition. Multigroup comparison by Kruskal-Wallis test and Tukey post-hoc test. Only the variables significant at a p-value < 0,05 are represented and ordered by increasing q-value (max q-value = 0,046), among 130 individual experiments. E. Correlation of PDL1 (x) with PVR, Nectin2, CD54, PDL2 and CD40 (y). « r » values are Spearman correlation coefficients.

Next, we analyzed the secretion of 32 DC derived cytokines and chemokines, and 17 T helper cytokines secreted by naïve CD4 T cells after 6 days of co-culture (Fig 3A, Table 5, Table 6). PDL1 high ICOSL low secreted the largest amount of most cytokines measured, such as TNFa, IL-1a, IL-1b, IL1RA, IL6, IL-10, IL12p40, IL-23, IL27, CCL19, BCA1, MIP1a, as compared to both PDL1 low ICOSL high DC and to Medium DC, but they did not induce more secretion of T helper cytokines by naïve CD4 T cells than Medium-DC (Fig 3B). Conversely, it was the PDL1 low ICOSL high DC that induced the highest activation of T cells as measured by the high expression of most CD4 T helper cytokines after co-culture, without a clear T helper polarization (Fig 3A, 3B, S3A, S3B). Therefore, we labeled PDL1 high ICOSL low DC the “secretory DC” and PDL1 low ICOSL DC the “helper DC”, both being different activated profiles, distinct from previously described tolerogenic DC. “Helper” DC increased very significantly the secretion of Th2 cytokines, IL-10, IL-3 and IL-9 by the CD4 T cells as compared to “secretory” DC, whereas IL-2 and IFNg were only mildly increased. There was no significant difference for Th17 cytokines.

**Fig 3.**
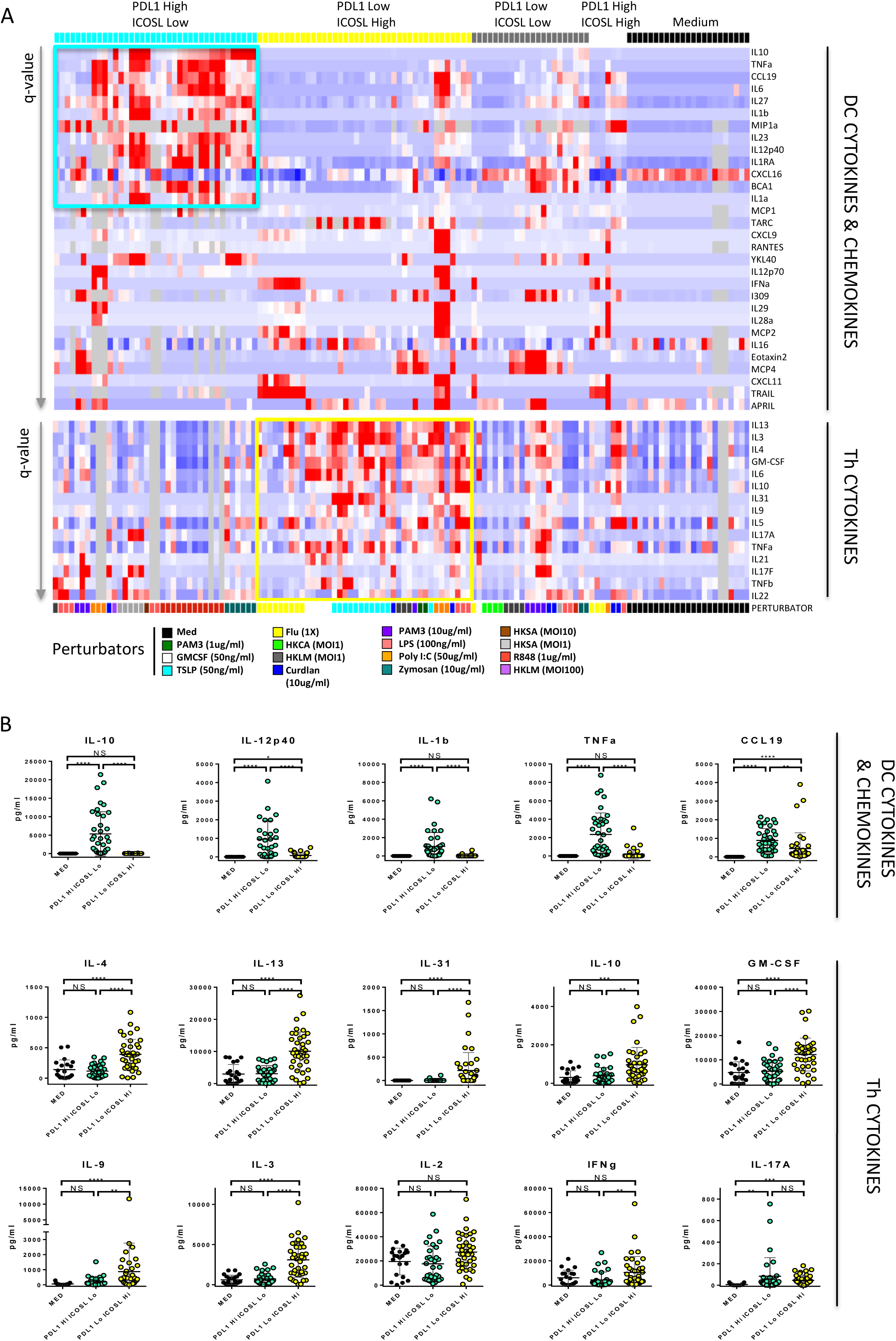
PDL1 and ICOSL expression pattern characterize “Secretory” and “Helper” DC. A. Heatmaps representing the cytokines and chemokines secreted by the DC measured in H24 supernatants (top), and the CD4 T helper cell cytokines measured after co-culture (bottom) in the 4 groups defined by PDL1 and ICOSL expression and Medium condition. Only the variables significant at a p-value < 0,05 after Kruskal-Wallis multigroup comparison and Tukey post-hoc test are represented and ordered by increasing q-value (max q-value = 0,035 (top) and 0,055 (bottom)), among 130 individual experiments. Cells in grey are missing values. B. Quantification of cytokines and chemokines secreted by the DC (top row) and of the CD4 T helper cell cytokines (2 bottom rows) in the Medium, PDL1 high ICOSL low and PDL1 low ICOSL high conditions.

To further characterize the changes occurring during DC activation in the context of cancer, we performed RNA sequencing of cDC2 sorted from HNSCC or blood and identified 882 differentially expressed genes (DEG): 639 increased in tumor cDC2 and 243 in blood cDC2 (Fig 4A, Table 7 for donors characteristics and Table 8 for DEG). Due to the minimal number of cells required for this experiment, inflamed tumors highly infiltrated by DC were necessarily selected (Fig S4A). In parallel, we compared transcriptomics data of cDC2 activated with pRNA, a TLR7/8 ligand expected to induce “secretory” DC or GM-CSF a “helper” DC2 inducer (Fig 4B) from GSE89442 (14). Using both comparisons of the stimuli together and towards unstimulated blood cDC2, we defined the “secretory” and “helper” signatures including 1473 and 1277 genes respectively (Fig 4C, Table 9). Among the 639 genes upregulated during tumor-induced maturation, 135 (21%) were shared with the “secretory” signature and only 64 (10%) with the “helper” signature, the 440 (69%) remaining genes being tumor-specific (Fig 4D). Using supervised lists of genes coding for checkpoints and maturation markers (Fig 4E left, Table 10), cytokines and chemokines (Fig 4E center, Tables 11 & 12), and of the NFkB pathway (Fig 4E right, Table 13), we confirmed that tumor cDC2 shared the majority of the genes with the pRNA “secretory” condition (Fig 4F). More in details, they overexpressed *CD274*/PDL1, and several other secretory specific markers identified previously at the protein level, such as *PDCD1LG2*/PDL2, PVR, *IL1B, IL12B, IL23A, TNF*, and *CCL19*, and also other negative checkpoints such as *IDO1, IDO2, and HAVCR2*/TIM3, and the migration marker *CCR7*.

**Fig4.**
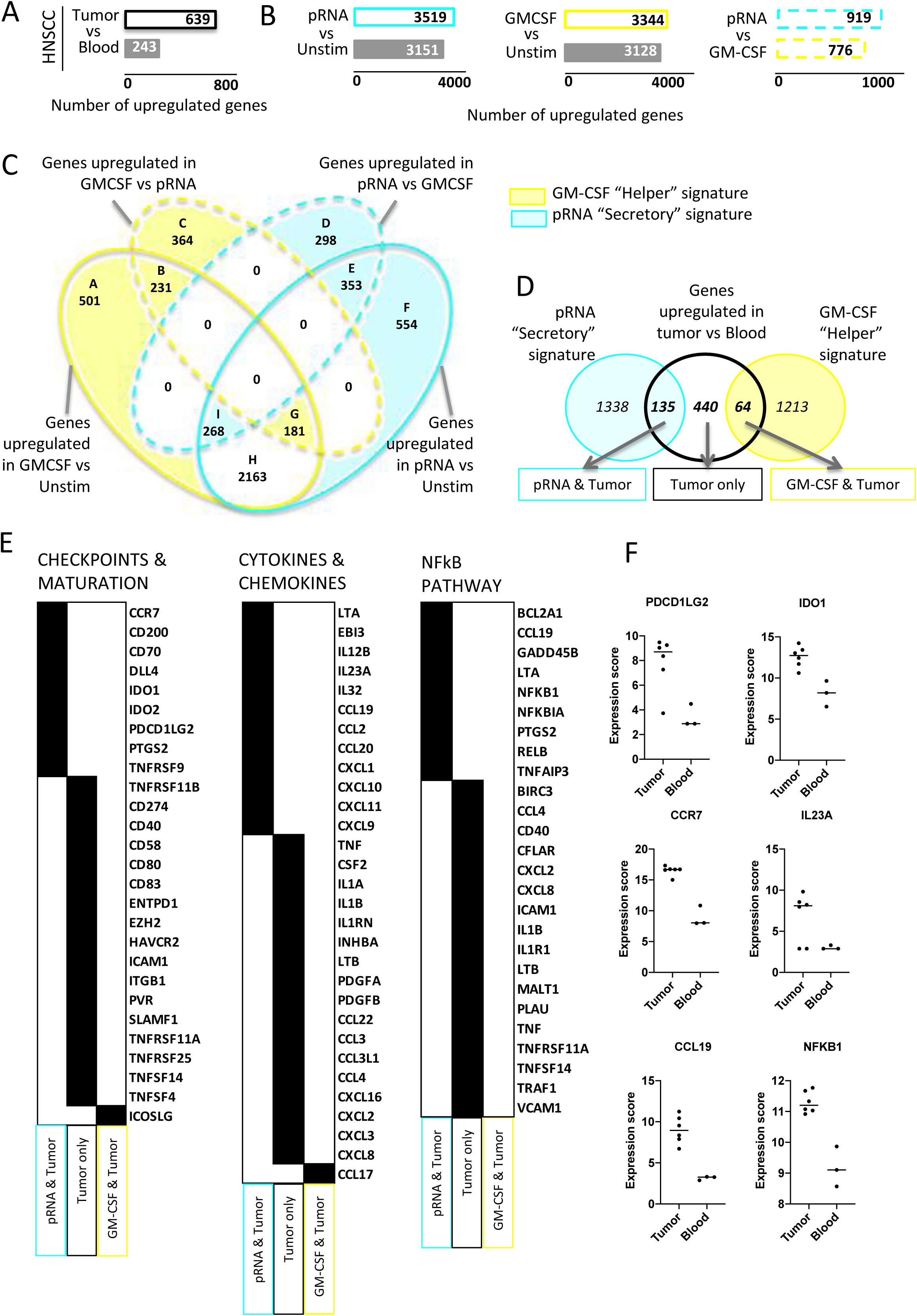
RNAseq of tumor vs blood cDC2 confirms that T cell-inflamed HNSCC are infiltrated by “secretory” DC. A. Analysis of differentially expressed genes (DEG) by DESeq2 between HNSCC tumor (n=6) and blood cDC2 (n=3). B. Analysis of DEG from dataset GS87442 by DESeq2 between unstimulated cell and pRNA, a TLR7/8 ligand (left) or GM-CSF (center) and pRNA vs GM-CSF (right). C. Venn diagram of upregulated genes identified in “B”. The blue and the yellow-colored area contain the genes of the “secretory” and “helper” signatures respectively. D. Venn diagram of the 639 tumor cDC2 upregulated genes with the “secretory” and “helper” signatures defined in “C”. E. Supervised analysis of the 135 genes shared between tumor & pRNA “secretory” signature (light blue), 440 tumor specific genes (black) and the 64 genes shared between tumor & GM-CSF (yellow), using 3 gene lists: checkpoint and maturation markers (left, 148 genes), cytokines and chemokines (center, 169 genes), NFkB pathway (right, 100 genes). F. Expression of selected genes in cDC2 from tumors and blood of HNSCC patients.

Since the concept of immature versus mature DC, and their respective roles in immune regulation, attempts have been made to identify classes of mature DC, such as “fully mature”, “immunogenic”, “inflammatory”, “semi-mature”, “tolerogenic” (3). These suffer from several limitations: 1) they lack a clear and consensual definition, 2) they lack universality and specificity, i.e many DC do not fall into any of these categories, or may fall into multiple. In this study, we report on a universal classification of human activated DC that mature either as “secretory” DC or as “helper DC”, recognizable by their opposed PDL1 and ICOSL expression. Each phenotype is induced by some specific stimuli, but not restricted to a single receptor pathway (Table 3). Tumor infiltrating cDC2 in inflamed HNSCC have the phenotypic signature of “secretory” DC. In blood and in non-inflamed HNSCC, DC have an immature phenotype (Fig 5). These observations have several applications for immunotherapies modulating DC in cancer and inflammatory diseases, such as DC stimuli used directly, or for DC-based vaccines, or even for standard cancer treatment that will increase danger signals in the tumor microenvironment. For example in cancer, the stimuli inducing “secretory DC” should be used in combination with anti-PD(L)1 antibodies, when it is not planned in some upcoming trials (NCT02320305, NCT03742804, NCT02180698), and the stimuli inducing “helper DC” could be used to increase the T cell response via polyfunctional Th cytokine profiles.

**Fig.5.**
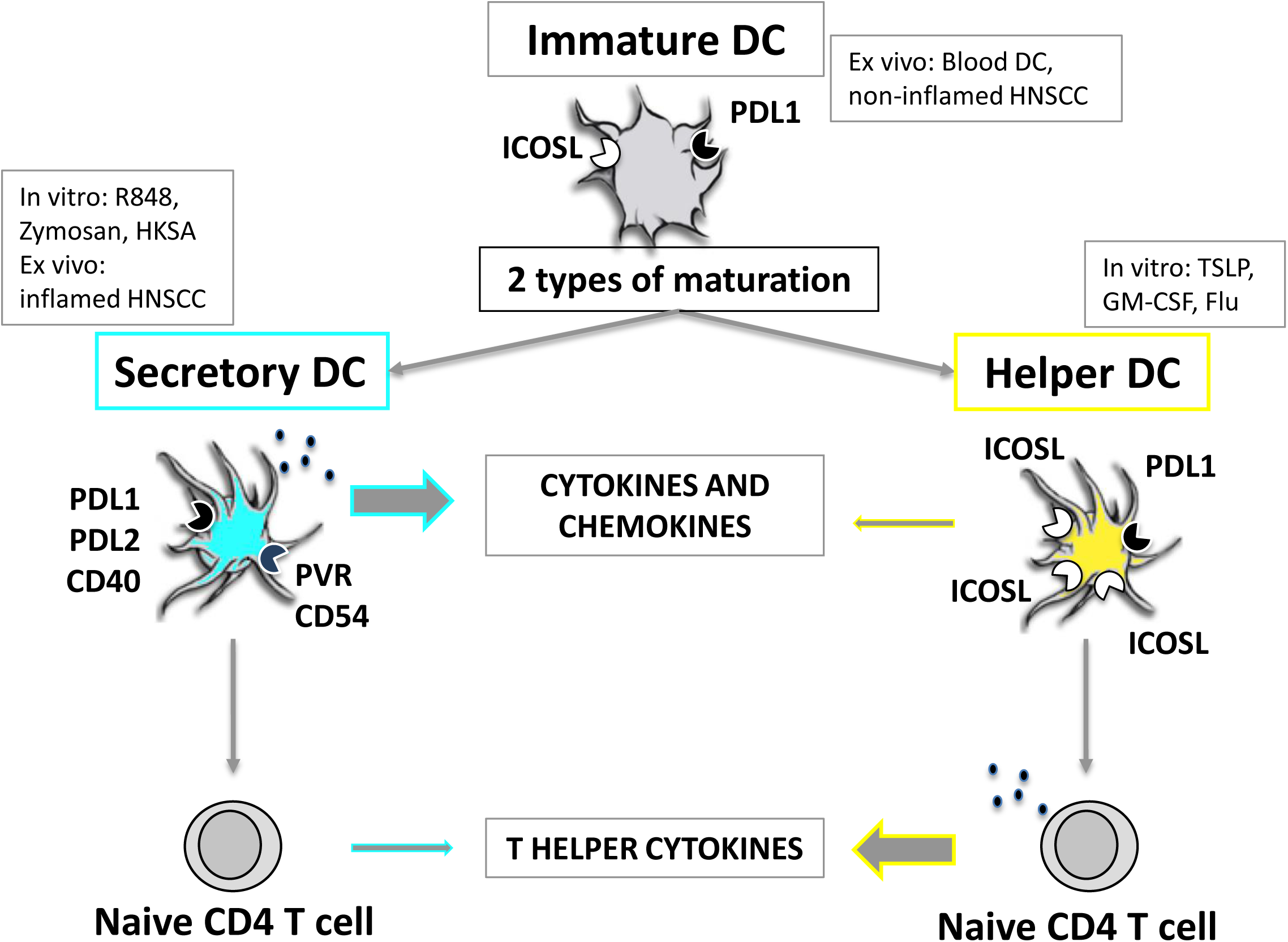
Schematic representation of DC activation into “helper” and “secretory” phenotypes.

## METHODS

### Human samples and patient characteristics

Fresh samples of HNSCC tumor tissues and blood of untreated patients with head and neck cancers were obtained from the pathology department of the Institut Curie hospital in accordance with the ethical guidelines, with the principles of Good Clinical Practice and the Declaration of Helsinki, and with patients consent. Patient characteristics for the flow cytometry cohort (Fig.1) and RNAseq cohort (Fig.3) are summarized in Supplementary Tables 2 and 7, respectively. Fourteen of 22 the patients of the FACS cohort were included in the clinical trial SCANDARE NCT03017573.

### Single-cell suspensions

Tumor tissues were mechanically and enzymatically digested in CO2-independent medium (Gibco) containing 5% FBS (HyClone). Enzymatic digestion consisted of three rounds of 15 min of incubation with agitation at 37 °C, separated by pipetting, with 2 mg/ml collagenase I (C0130, Sigma), 2 mg/ml hyaluronidase (H3506, Sigma) and 25 µg/ml DNAse (Roche). The samples were filtered on a 40-µm cell strainer (Fischer Scientific) and were diluted in PBS 1X (Gibco) supplemented with EDTA 2 mM (Gibco) and 1% de-complemented human serum (BioWest). After centrifugation, cells were suspended in the same medium and were counted by trypan blue before being assessed by flow cytometry or sorted. PBMC were isolated from blood samples using FICOLL (GE Healthcare) gradient centrifugation.

### Antibodies, flow cytometry and cell sorting

Single-cell suspensions from digested tumor and from blood were stained with antibodies (Table 14) for 15 min at 4°C. After washing step, cells were analyzed or sorted directly, immediately after having added DAPI (Miltenyi Biotec) for dead cells exclusion. Flow cytometry phenotyping was performed on BD LSRFortessa Analyzer. Cell sorting for the RNA-seq experiment were performed on BD FACSAria III using the purity and low-pressure mode, and a 100-μm nozzle. DC subsets and MMAC were sorted in Eppendorf tubes containing TCL buffer (Qiagen) supplemented with 1% β–mercaptoethanol (SIGMA) before RNA extraction, as decribed in Michea P, Noël F et al. (15).

### In vitro analysis

Material and methods are described in details in the resource paper from Grandclaudon et al. (13). As compared to the resource paper containing 118 data points for primary blood CD11c+HLA-DR+ cells (referred to as bDC), we generated supplementary experiments and analysis to specifically address our question. We added 36 data points for the analysis of surface markers (leading to a total of 154 data points) among which 12 for the analysis of DC secreted cytokines and chemokines and of the T helper cytokines (leading to a total of 130 data points). Extra data points included: Curdlan 10ug/ml (n=1), Flu (1X) (n=3), Flu(1X)+TSLP(50ng/ml) (n=3), HKSA (MOI10) (n=3), GM-CSF 50ng/ml (n=4), LPS (n=3), Medium (n=9), Poly I:C 50ug/ml (n=4), R848 1ug/ml (n=3), TSLP 50ng/ml (n=3), for a total of 29 blood donors. The antibodies used for the checkpoints and maturation markers analyzed by flow cytometry are listed in Table 4. For the DC secreted cytokines and chemokines, we measured 24 supplementary cytokines and chemokines. IL1a, IL1b, IL6, IL10, TNFa and IL12p70 were measured by cytometry bead assay flex set (CBA) and we added the measure of IFNa. IL23 and IL28a were measured by Luminex and we added the measure of APRIL, BCA1, CCL19, CXCL11, CXCL16, CXCL9, Eotaxin2, I309, IFNb, IL12p40, IL16, IL1RA, IL27, IL29, IP10, MCP1, MCP2, MCP4, MIP1a, RANTES, TARC, TRAIL, YKL40 (Table S5). The 17 T helper cytokines were analyzed by CBA or Luminex (Millipore) (Table 6), similarly to the resource paper.

### RNA extraction, sequencing and data pre-processing

Material and methods are described in details in the resource paper (15). Briefly, single Cell RNA Purification Kit (Norgen Bioteck) was used for RNA extraction, including on-column DNase digestion (Qiagen), as described by the manufacturer’s protocol. RNA integrity was controlled with a RNA 6000 Pico Kit (Agilent Technologies) in BioAnalyzer. cDNA was generated with SMARTer Ultra Low input RNA for Illumina Sequencing-HV (Clontech), following manufacturer’s protocol with 14 cycles for amplification. Quality controls were performed with Qubit dsDNA high sensitivity (Thermofisher) and an Agilent Bioanalyzer using nanochip (Agilent Technologies). Multiplexed pair-end libraries 50nt in length were obtained using Nextera XT kit (Clontech). Sequencing was performed in a single batch with Illumina HiSeq 2500 using an average depth of 15 million reads. Library, sequencing and quality controls were performed by the NGS facility at the Institut Curie. Reads were mapped to the human genome reference (hg19/GRCh37) using Tophat2 version 2.0.14. Gene expression values were quantified as read counts using HTSeq-count version 0.6.1. Genes with less than one read count in at least one sample were filtered out and. The remaining raw data were normalized and analyzed using DESeq2 R package. Differentially expressed genes were obtained with an adjusted p-value of 0,10. The supervised list of genes used in Fig 4D were established by including all markers analyzed at the protein level in the in vitro analysis and by adding other known checkpoints and maturation markers, cytokines and chemokines from literature search. The NFkB pathway genes list was established by literature search.

#### Data availability

RNA-seq data that support the findings of this study will been deposited in the NCBI Sequence Read Archive (SRA).

### Analysis of Flow cytometry data

We measured a total of 434 parameters including 52 cell/cell ratios. We established a sub-list of 81 non-redundant parameters, meaning that each population was expressed in percentage of its parental population. The list of 81 parameters was used in Fig 1D, and the list of 434 parameters enriched wit 14 clinical parameters was used for the elastic net model in Fig S1D. The elastic net model was performed using R software, a Lambda at 1SE, and an alpha of 0,5.

### Statistical analysis

Statistical analyses of flow cytometry data (Fig1) and in vitro analysis (Fig2, Fig3) were performed using ANOVA or Kruskal-Wallis tests for parametric and non-parametric data respectively, with Qlucore and GraphPad Prism 8 (GraphPad Software Inc.) softwares. Data were considered significant for adjusted p-values after Tukey or Dunn’s tests superior to 0.05. t-SNE was performed using Qlucore software and a perplexity of 15.

## ACKNOWLEDGMENTS

The authors wish to thank the INSERM U932; the Institut Curie Flow-Cytometry facility, in particular Zofia Maciorowsky, Annick Viguier, and Sophie Grondin for their technical help and expertise; the Institut Curie NGS platform; François Lemoine, Géraldine Lescaille, and Chloé Bertolus at Hôpital Pitié Salpêtrière, Paris, France, for providing tissues and blood from 4 patients, samples further used for transcriptomics analysis. This study makes use of data generated by Mathan et al., from the Radboud Institute for Molecular Life Sciences, Nijmegen, Netherlands, identified by the Gene Expression Omnibus GSE89442.

## SUPPLEMENTARY FIGURES LEGEND

**FigS1A**. T cell panel gating strategy

**FigS1B**. Left: Flow cytometry staining for BDCA3 expression in 4 cell populations in a HNSCC primary tumor. This tumor was selected for its high level of cDC1 infiltration. Right: Percentages of cDC1, gated as BDCA3 high, in the cDC1e gate (n = 6 tumors).

**Fig S1C**. Flow cytometry staining showing CD15 expression in Lin-HLADR-population. Most Lin-HLADR-CD11c+ cells are CD15+, therefore having a neutrophil phenotype.

**Fig S1D**. Heatmap representing the expression of PDL1 and ICOSL in the 4 subsets of CD11c+HLA-DR+ cells in the 22 HNSCC samples, ordered by the level of CD3 infiltration from the lowest (left) to the highest (right).

**Fig S1E**. Elastic net model of the 434 parameters measured by flow cytometry and 14 clinical parameters in the 22 HNSCC, showing the parameters the most representative of CD3 Low, CD3 Int and CD3 High tumors. The “Live” gate was established by selecting the live cells among a parental gate of all the cells in the FSC-A versus SCC-A graph, excluding only the debris and red blood cells. The “Live Lymphocyte” gate was established by selecting the live cells among a parental gate of cells having the FSC and SCC levels corresponding to lymphocytes only.

**Fig S3A**. Quantification of cytokines and chemokines secreted by the DC, in the Medium, PDL1 high ICOSL low and PDL1 low ICOSL high conditions.

**Fig S3B**. Quantification of the CD4 T helper cell cytokines, in the Medium, PDL1 high ICOSL low and PDL1 low ICOSL high conditions.

**Fig S3C**. T cell expansion at day 6 of DC-T co-culture in the Medium, PDL1 high ICOSL low and PDL1 low ICOSL high conditions.

**FigS4A**. Flow cytometry sorting strategy for RNA sequencing of blood and tumor infiltrating cDC2, selected as CD45+, CD3-, CD19-, CD56-, CD11c+, HLA-DR+, CD14-, BDCA1+. Plots from a representative donor.

**ALL TABLES ARE PROVIDED AS SUPPLEMENTARY MATERIAL**

