## Supplementary figures and images for "Pdl1 and icosl discriminate human secretory and helper dendritic cells"

### Supplementary Figures

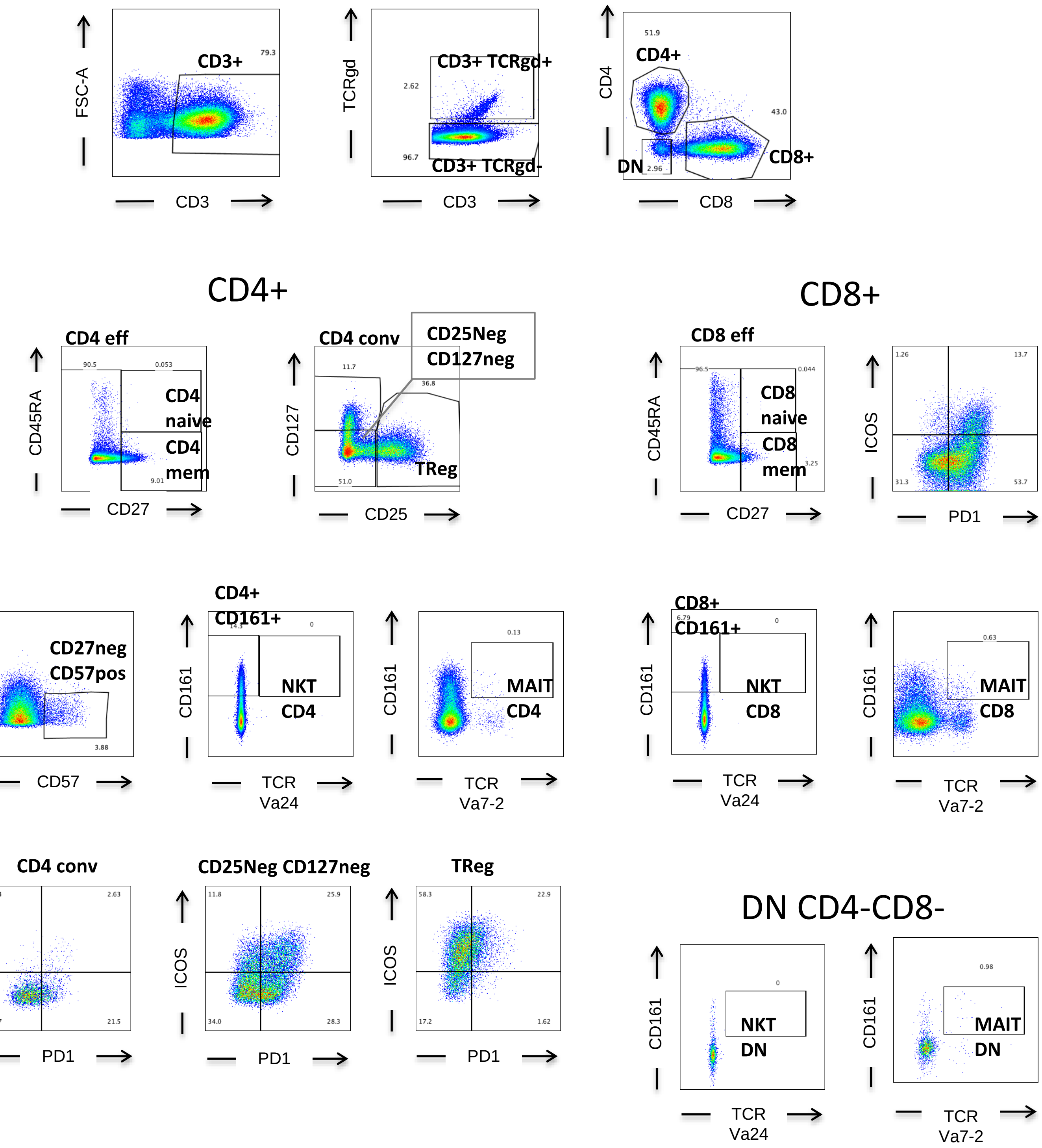

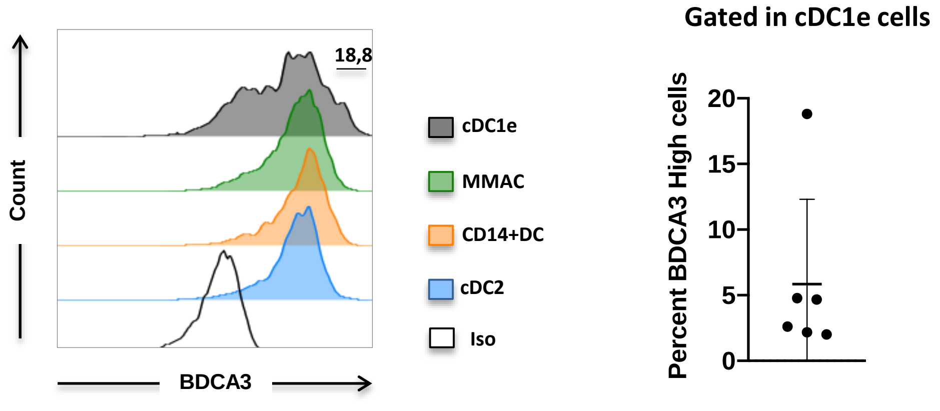

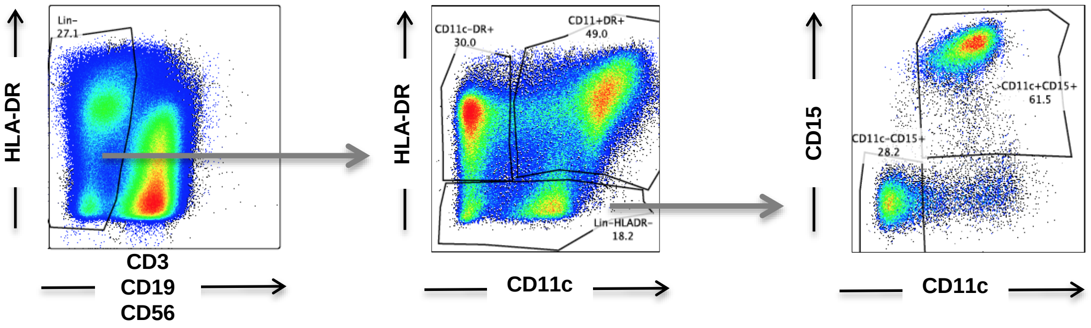

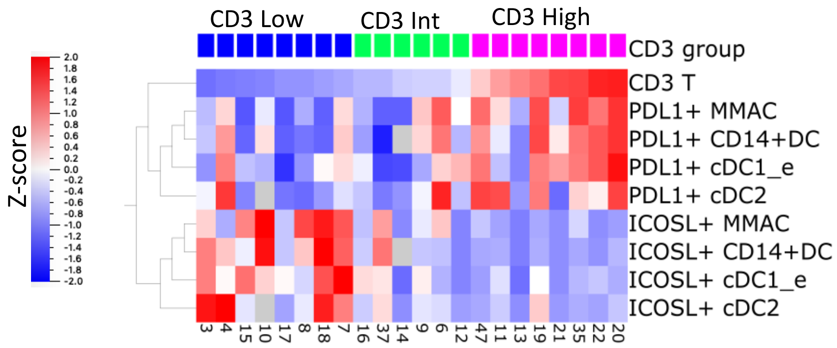

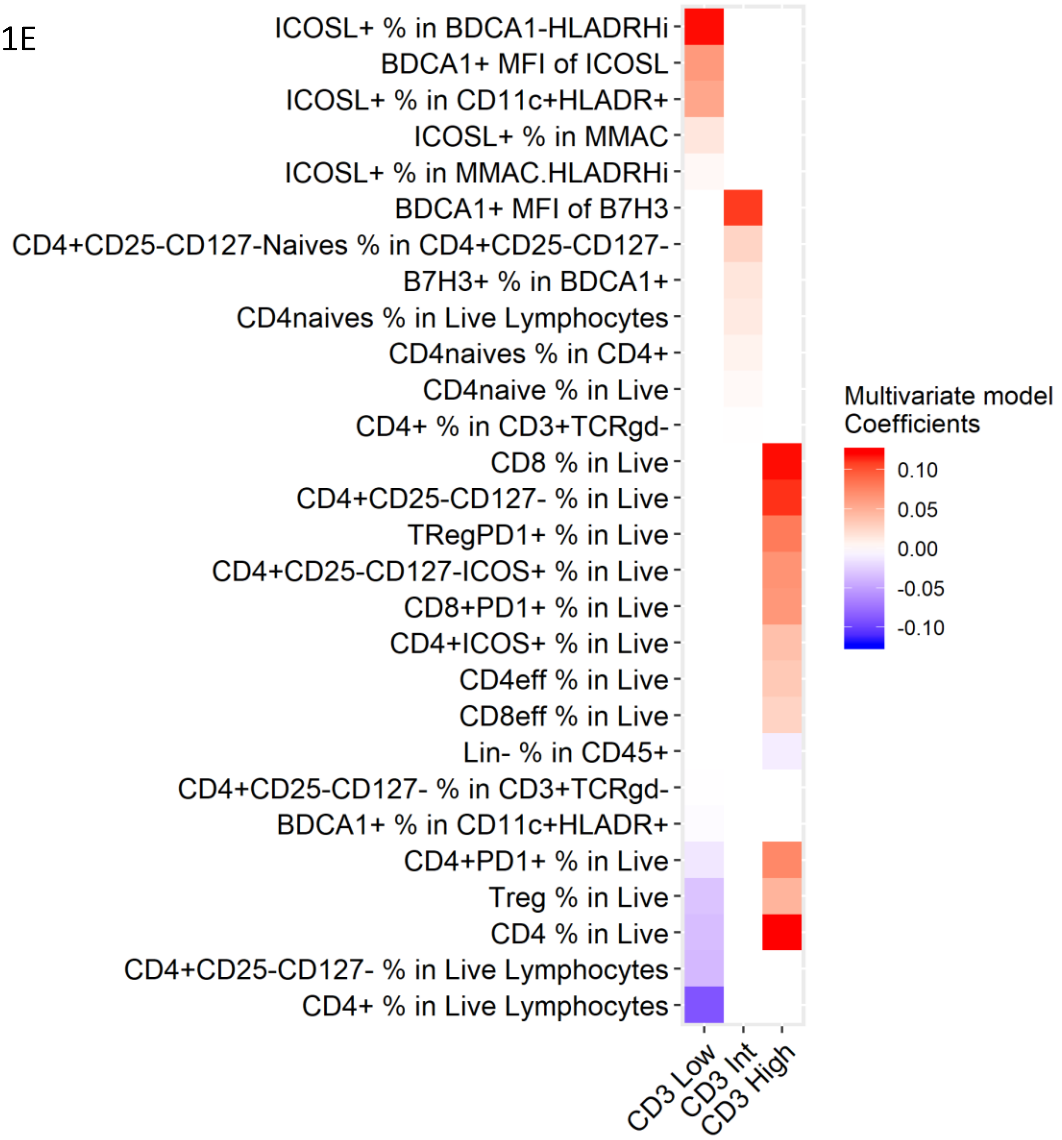

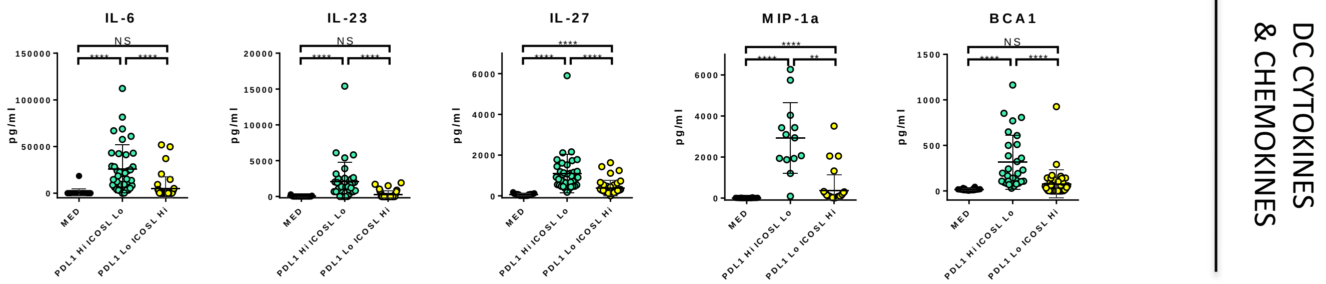

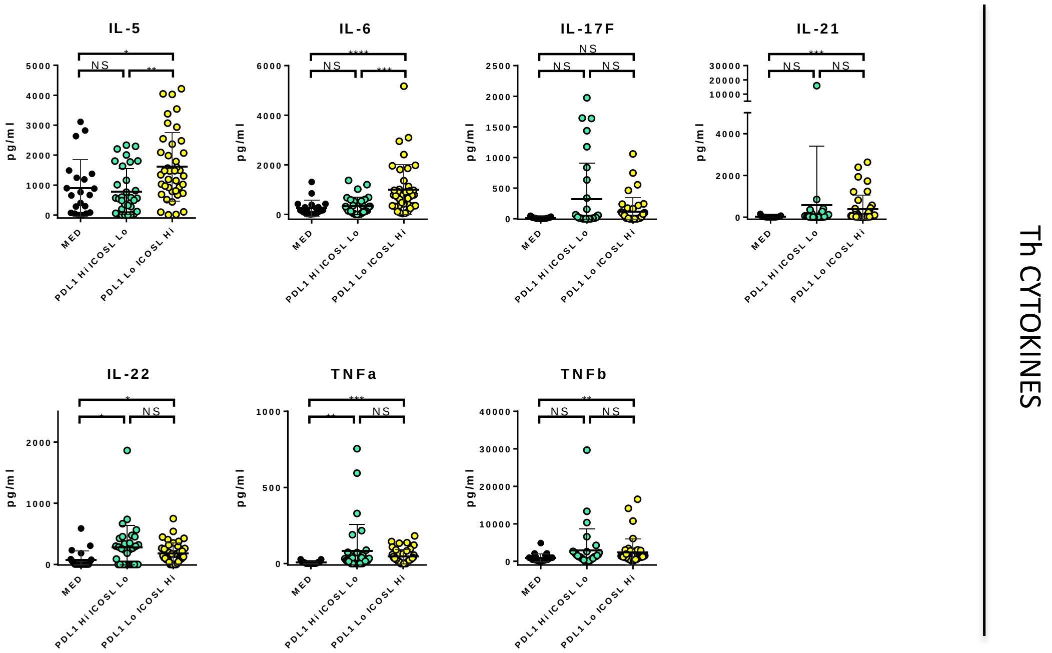

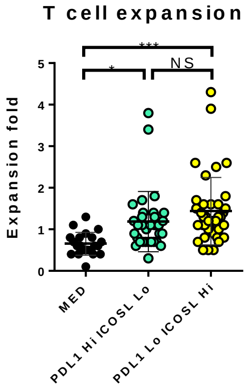

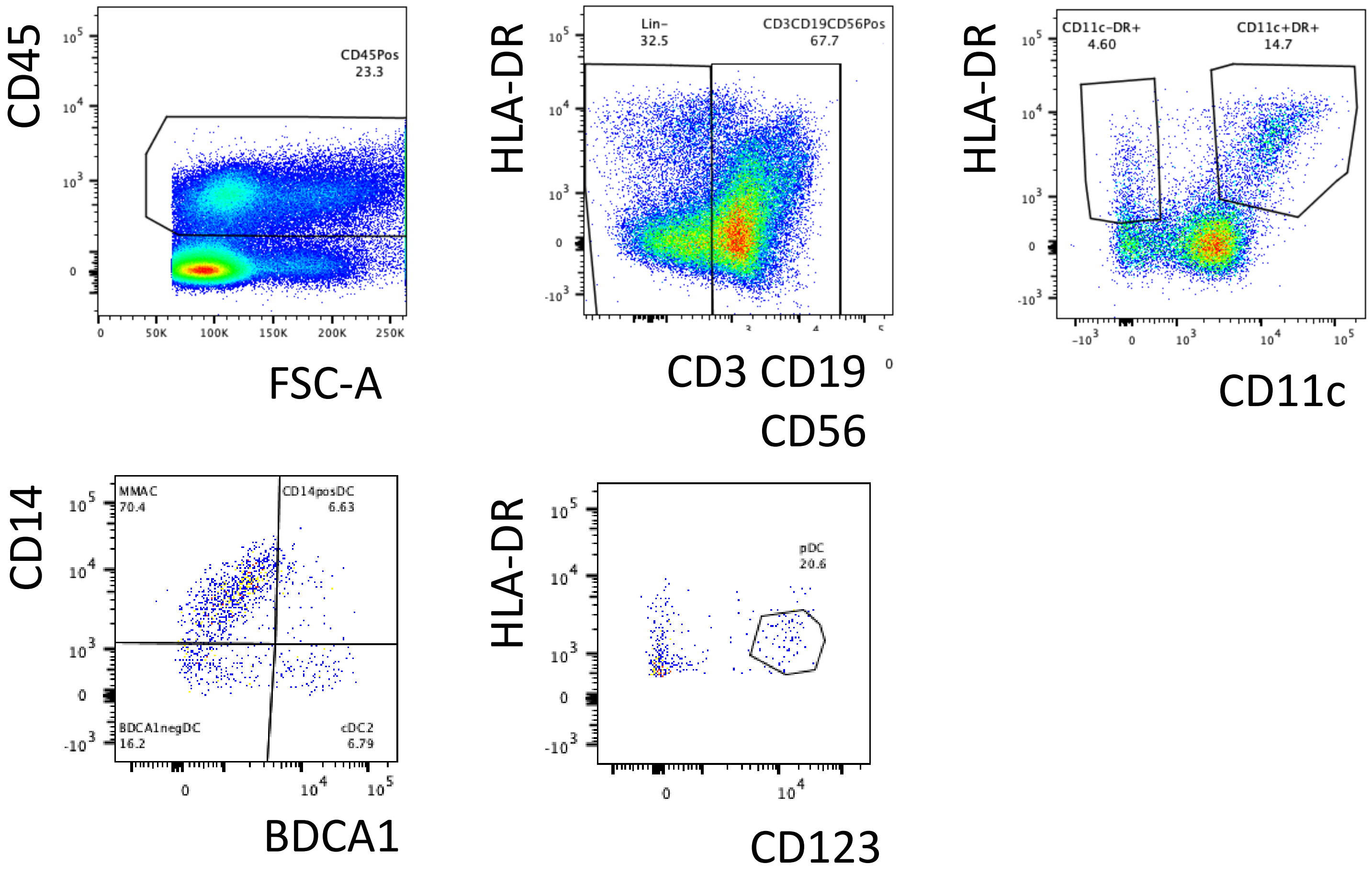
